# Range-wide population genetics reveals how disjunctions, admixture, and translocations influence conservation of *Physaria globosa* (Brassicaceae)

**DOI:** 10.64898/2026.02.05.703860

**Authors:** Wen-Hsi Kuo, Clara Landon, Burgund Bassüner, Alexander G. Linan, Tara R. Littlefield, Matthew A. Albrecht, Christine E. Edwards

## Abstract

Population genetic analyses of endangered species can directly inform management strategies to conserve genetic diversity and increase long-term survival. *Physaria globosa*, an endangered mustard, has a geographically restricted and disjunct distribution across river systems in Indiana, Kentucky, and Tennessee, USA. In this study, we assessed whether genetic structure was shaped by the species’ disjunct distribution, whether genetic diversity has been affected by bottlenecks, drift, or inbreeding, and whether translocations based on mixed sources can retain genetic variation. We genotyped 332 individuals at 20 microsatellite loci from 14 natural and 2 translocated populations from across the species’ range. The genetic structure of natural populations showed two major geographic groups with little to no connectivity between them: a northern group in Indiana and Kentucky (IN-KY) and a southern group in Tennessee (TNE-TNW). Genetic diversity was highly variable among and within geographic regions; however, several local populations showed reduced genetic diversity, inbreeding, or recent bottlenecks, consistent with demographic decline and ongoing pressures from habitat degradation and competition from invasive plants. The two translocated populations showed clear genetic mixing of source populations but showed depletion of private alleles. Our results suggest that conservation actions for *P. globosa* should prioritize the protection of multiple populations within each major geographic region. Combining material from mixed sources appears to be a viable approach to increase genetic diversity and restore gene flow, but we recommend that translocations combine material from within the same geographic regions to preserve the distinct evolutionary histories of the main geographic groups.

## Introduction

Genetic diversity is widely recognized as the raw material underlying phenotypic and functional variation within species, and its maintenance is a central goal of biodiversity conservation (Shaw et al. 2025). Greater genetic diversity is often associated with greater resilience to environmental change, and increased evolutionary potential and individual fitness. Genetic diversity can also improve the establishment and long-term persistence of reintroduced populations (Hoelzel, Bruford, & Fleischer 2019; Van Rossum & Le Pajolec 2021). For endangered species with narrow distributions and ongoing habitat loss and degradation, conserving genetic diversity requires consideration of natural populations, *ex situ* collections, and conservation translocations derived from them (Aguilar-Gómez et al. 2025; Edwards et al. 2025; Linan et al. 2024; Mashburn et al. 2023; Van Rossum et al. 2025).

Conservation strategies for rare plant species generally span a continuum from protecting extant natural populations to more interventionist approaches involving the establishment of *ex situ* populations and translocations into suitable habitats (Bucharova 2017; Gaitán-Espitia & Hobday 2021). In plants, these interventions can include controlled crosses among natural populations and translocation of individuals among populations to promote interbreeding and restore genetic variation (Edwards et al. 2025). Such actions can be valuable when fragmentation, demographic decline, or habitat degradation have reduced effective population sizes and limited natural gene flow. In these circumstances, facilitated gene flow can increase genetic diversity, reduce inbreeding, and potentially enhance fitness and longer-term adaptive potential. However, outcomes are often context-dependent and not always predictable; thus, their application remains debated (MacDonald et al. 2025).

Using mixed-source materials for controlled crosses, population augmentations, translocations, and other forms of assisted migration raise several important questions in conservation genetics. Do disjunct populations represent distinct historical or adaptive components that should be protected separately (Fraser & Bernatchez 2001)? Under what conditions does the risk of loss of genetic diversity due to inbreeding or genetic drift in an isolated population exceed the risk associated with admixture? Can mixed populations capture sufficient genetic diversity to maintain their persistence in local or changing environments? The major concerns are well known: introduced gene flow may cause outbreeding depression, disrupt local adaptation, generate maladapted genotypes, or generate ecological mismatches in recipient environments (Bucharova 2017; Hufford & Mazer 2003; MacDonald et al. 2025). Hybridization can also lead to genetic or demographic swamping if rare lineages are overwhelmed by introgression from more widespread relatives (Aguilar-Gómez et al. 2025). In addition, small, inbred but viable populations may have already purged some deleterious recessive alleles, such that genetic rescue could introduce new genetic load and reduce fitness (Dussex, Morales, Grossen, Dalen, & van Oosterhout 2023). These risks do not preclude intervention but underscore the need to identify a clear biological and demographic rationale, carefully evaluate risks of both source and recipient populations, and conduct post-release genetic monitoring to assess outcomes (Aguilar-Gómez et al. 2025; Linan et al. 2024).

Globe bladderpod, *Physaria globosa* (Desv.) O’Kane & Al-Shehbaz, is a federally endangered annual to biennial herbaceous mustard with a geographically restricted and discontinuous distribution in the southeastern United States (Fuentes-Soriano & Kellogg 2021). It occurs on steep, rocky bluffs and talus slopes, usually on calcareous substrates with south-to west-facing slopes near rivers or streams (Fig. 1A). The species is distributed across four disjunct regions: the confluence of the Wabash and Ohio Rivers in Indiana (IN), the Kentucky River watershed in Kentucky (KY), and the Cumberland River watershed in Tennessee (TN), which is separated into eastern (TNE) and western (TNW) groups, divided by the Nashville metropolitan area (Fig. 2). *Physaria globosa* is threatened by habitat loss and degradation, soil erosion, and woody encroachment, particularly from invasive nonnative shrubs (Kallison et al. 2026; Long, Albrecht, Call, & Region 2021). Despite its conservation status, it remains unclear how genetic variation is distributed across these disjunct regions and whether individual populations are experiencing genetic risks such as bottlenecks, inbreeding, or genetic erosion.

**Figure 1.**
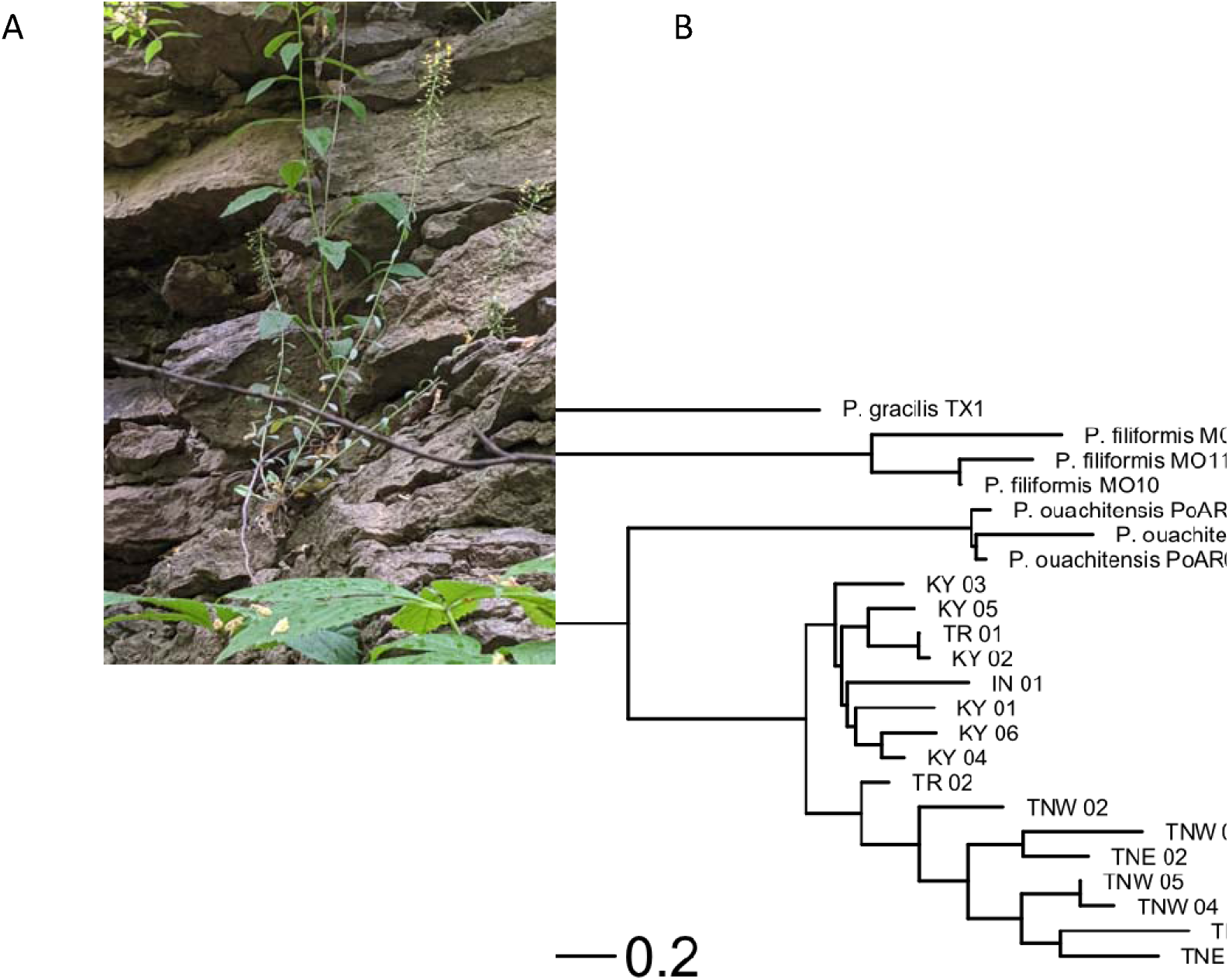
Photos of *Physaria globosa* and genetic distance tree. (**A**) Native habitat. (**B**) Neighbor-Joining tree based on Nei’s genetic distances. The tree was rooted at *P. gracilis*.

**Figure 2.**
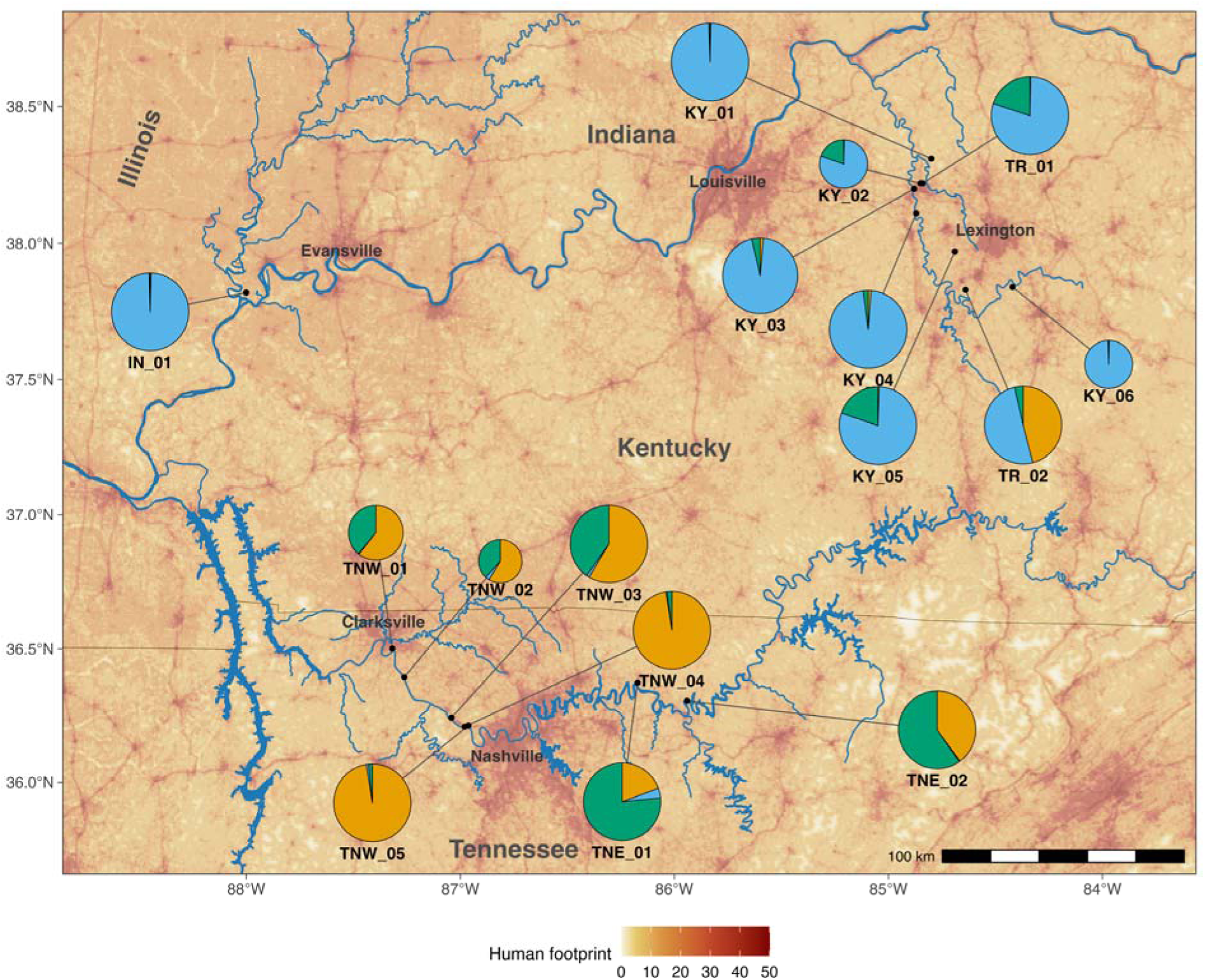
Map of *P. globosa* collection localities, with population structure K = 3 shown in pie charts. The size of the pie chart reflects the sampling size per population. See **Table 1** for detailed information.

**Table 1.**
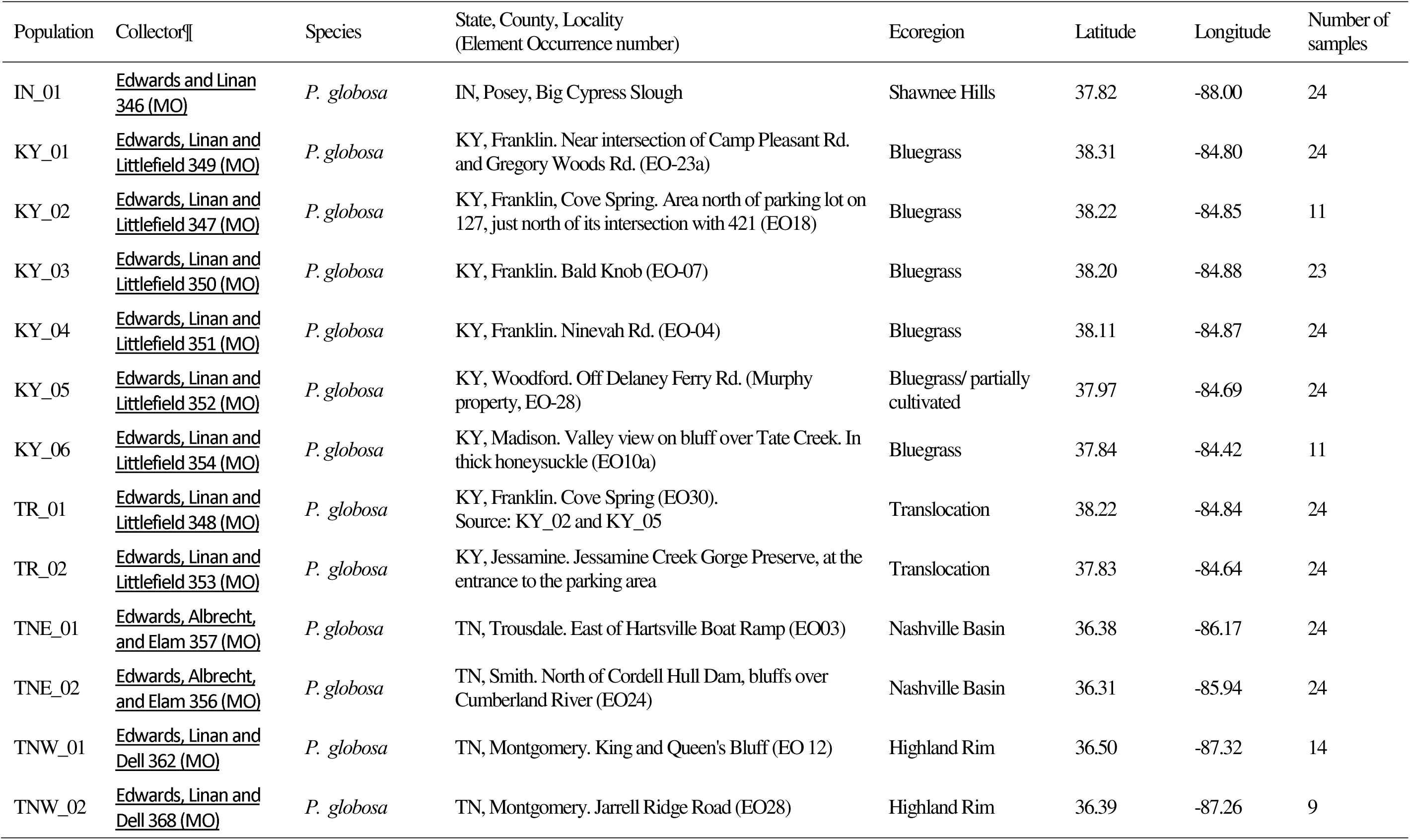
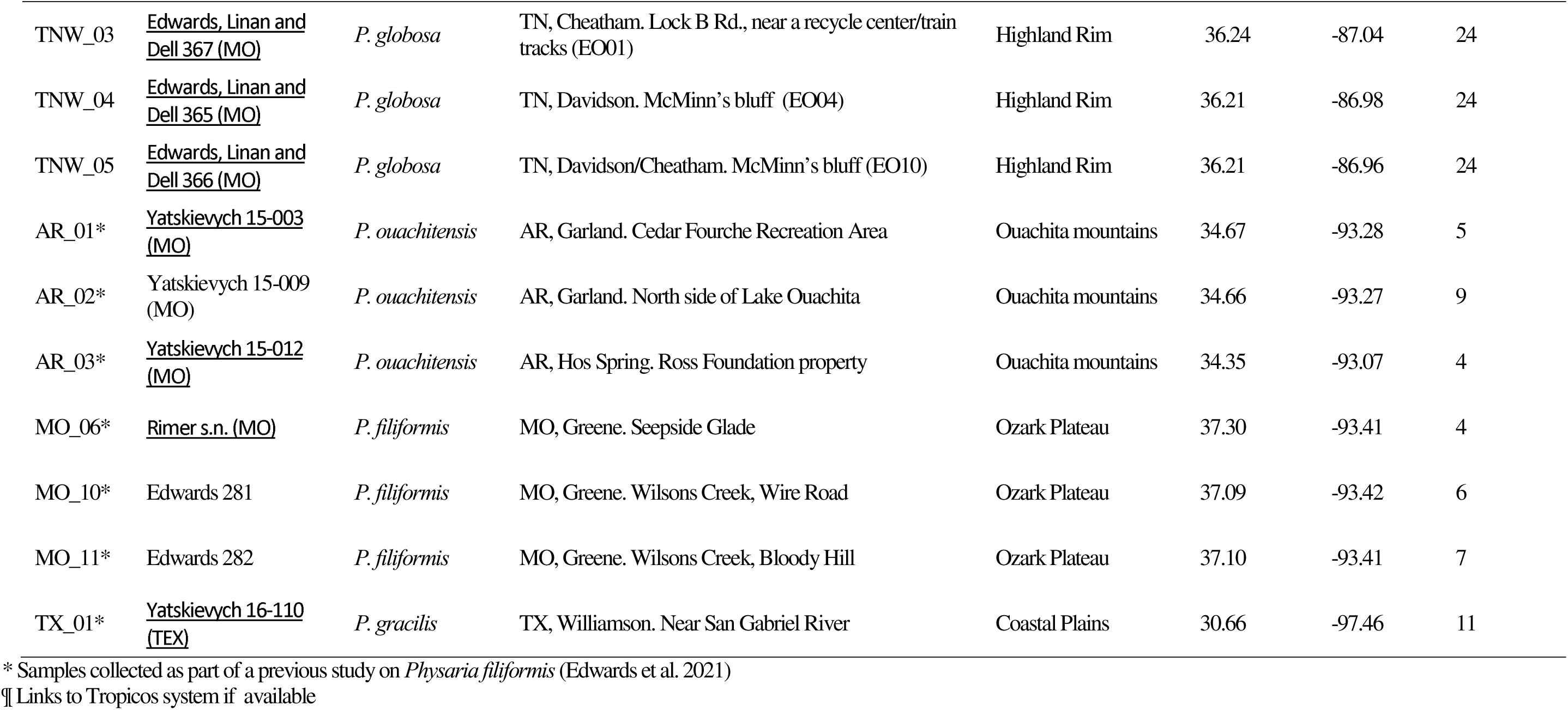
Information about the populations and samples of *Physaria globosa* included in the study, including voucher number, collection locality, ecoregion, GPS coordinates, and number of samples collected per site.

Climate change may further intensify these threats, as extended summer droughts and extreme precipitation events can increase both physiological stress and flood-or erosion-related habitat disturbance (Donat, Lowry, Alexander, O’Gorman, & Maher 2016; Xu et al. 2019). Together, these ecological and genetic vulnerabilities make *P. globosa* a useful system for evaluating whether conservation efforts should prioritize conserving existing population structure in natural populations and *ex situ* conservation collections or restoring genetic diversity through translocation using mixed-source material to facilitate admixture. Although such interventions may help maintain genetic diversity and promote long-term persistence, they also carry risks, including outbreeding depression, disruption of adaptive differentiation, and incomplete representation of natural genetic diversity in *ex situ* collections. Management decisions should therefore be guided by range-wide data on genetic diversity and population structure, while also evaluating the consequences of using multiple sources for conservation translocations.

This study addresses these questions using a comprehensive range-wide analysis of genetic diversity and structure in *P. globosa*. We sampled 14 natural populations spanning the species’ distribution and two mixed-source translocated populations. We genotyped 332 individuals at 20 microsatellite loci. Our goals were to understand: 1) whether genetic structure was shaped by the species’ disjunct distribution, 2) whether genetic diversity in populations has been affected by bottlenecks, drift, or inbreeding; and 3) whether existing mixed-source translocations retain genetic variation while remaining consistent with the broader recovery goal of promoting representation, resiliency, and redundancy across the range of this endangered species. Finally, we evaluated the conservation implications of these patterns for both *in situ* protection and *ex situ* management.

## Materials and Methods

### Study species

*Physaria globosa* is a member of the mustard family (Brassicaceae). It is morphologically distinct from other eastern and south-central U.S. congeners, including *P. ouachitensis* and *P. filiformis*, which are annual plants with fibrous or slender root systems, whereas *P. globosa* is most frequently biennial or sometimes annual, and often develops a woody caudex (Flora of North America, *Flora of North America* 2010; Fuentes-Soriano & Al-Shehbaz 2013; Fuentes-Soriano & Kellogg 2021; Kallison et al. 2026). Plants may produce multiple stems and grow up to 50 cm tall. Small yellow flowers are produced in April and May and are pollinated primarily by small native bees and flies (Krosnick et al. 2022). In June and July, the species produces distinctive globular fruits that give bladderpods their common name. Seeds are thought to be dispersed by water and gravity along the rivers (Long et al. 2021), or transported by small mammals (Tara Littlefield, pers. observation). We have observed that *Physaria globosa* seeds can persist in soil for at least five years (unpublished data), which is common in the genus *Physaria* (Elberling 2000).

### Sample collection and DNA extraction

We sampled *Physaria globosa* across its known geographic range in May 2021, with sampling effort in each region proportional to the number of known occurrences in each of the four main geographic regions (**Table 1**). In total, we collected 332 individuals from 16 populations. These included the only known natural population in Indiana, USA (IN_01); seven natural populations in Tennessee, USA, including two eastern Tennessee populations (TNE_01, TNE_02) and five western Tennessee populations (TNW_01, TNW_02, TNW_03, TNW_04, TNW_05); six natural populations in Kentucky, USA (KY_01, KY_02, KY_03, KY_04, KY_05, KY_06); and two translocated populations in Kentucky, USA (TR_01, TR_02).

The two translocated populations had distinct establishment histories. TR_01 was established by Tara Littlefield using two sources of introduced material. The first source was from a road-cut rescue at Cove Spring, Kentucky, USA (KY_02) in 2014. Approximately 100 seeds from 30-40 maternal plants were collected and moved to a suitable habitat on the east end of Cove Spring Park, approximately one mile away from the original site. The introduction site had recently been cleared of honeysuckle prior to seed addition, within an approximately 5 × 5 m managed area. The habitat included limestone rock outcrops and associated woody species such as *Juniperus virginiana*, *Quercus muehlenbergii*, *Fraxinus quadrangulata*, *Cercis canadensis*, and *Ostrya virginiana*. The second source consisted of seeds collected from Woodford County, Kentucky, USA (KY_05) in 2015. Approximately 200 additional seeds from 5-6 maternal plants were introduced to the site. Honeysuckle removal continued in winter 2015, with management focused on expanding the original cleared area where *Physaria globosa* was introduced. Therefore, the TR_01 population could be described as a field-established introduction with mixed seed-source origins. TR_01 has been monitored annually by Tara Littlefield, local volunteers, and staff from the Office of Kentucky Nature Preserves. No plants were observed inside the original 5 × 5 m managed area in 2014, followed by two vegetative plants in 2015, six vegetative and one flowering plant in 2016, with continued annual increases in flowering plants through 2025. Later increases appear to coincide with continued habitat management and honeysuckle removal at the site. The TR_01 population reached 238 vegetative and 29 flowering plants in 2025.

The TR_02 population at Jessamine Creek Gorge Preserve was established by a local conservation enthusiast. To the best of our knowledge, plants were originally sampled from populations in both Kentucky and Tennessee in the early 1990s and established on private property in a mixed-source population. These plants were allowed to naturally cross-pollinate. Seeds resulting from the mixed-source population were harvested and were broadcast-sown approximately three times at Jessamine Creek Gorge Preserve from 2010 to 2020. The specific source locations for the original population and the number of seeds used in the translocation were not recorded.

For both wild and translocated populations, we sampled leaf tissue from 10 -24 individuals per population, depending on population size and site accessibility. We also collected one herbarium voucher specimen per population, which was deposited in the Missouri Botanical Garden Herbarium (MO). The sampling locations, adjacent rivers, main cities, and human footprint (Gassert et al. 2023) were plotted in **Fig. 1**. To provide genus-level context for genetic diversity estimates, we also included 17 individuals from three populations of *P. filiformis* (Rollins) O’kane and Al-Shehbaz in Missouri, USA, 19 individuals from three populations of *P. ouachitensis* Kallison and C.E. Edwards in the Ouachita Mountains of Arkansas, USA, and 12 individuals from one population of *P. gracilis* (Hook.) O’kane and Al-Shehbaz in Texas, USA (**Table 2**), all previously included in C. E. Edwards et al. (2021). The final dataset comprised 406 individuals. Leaf tissue was dried in silica gel in the field, and DNA was extracted from silica-dried tissue using a modified CTAB protocol (Allen, Flores-Vergara, Krasynanski, Kumar, & Thompson 2006).

**Table 2.**
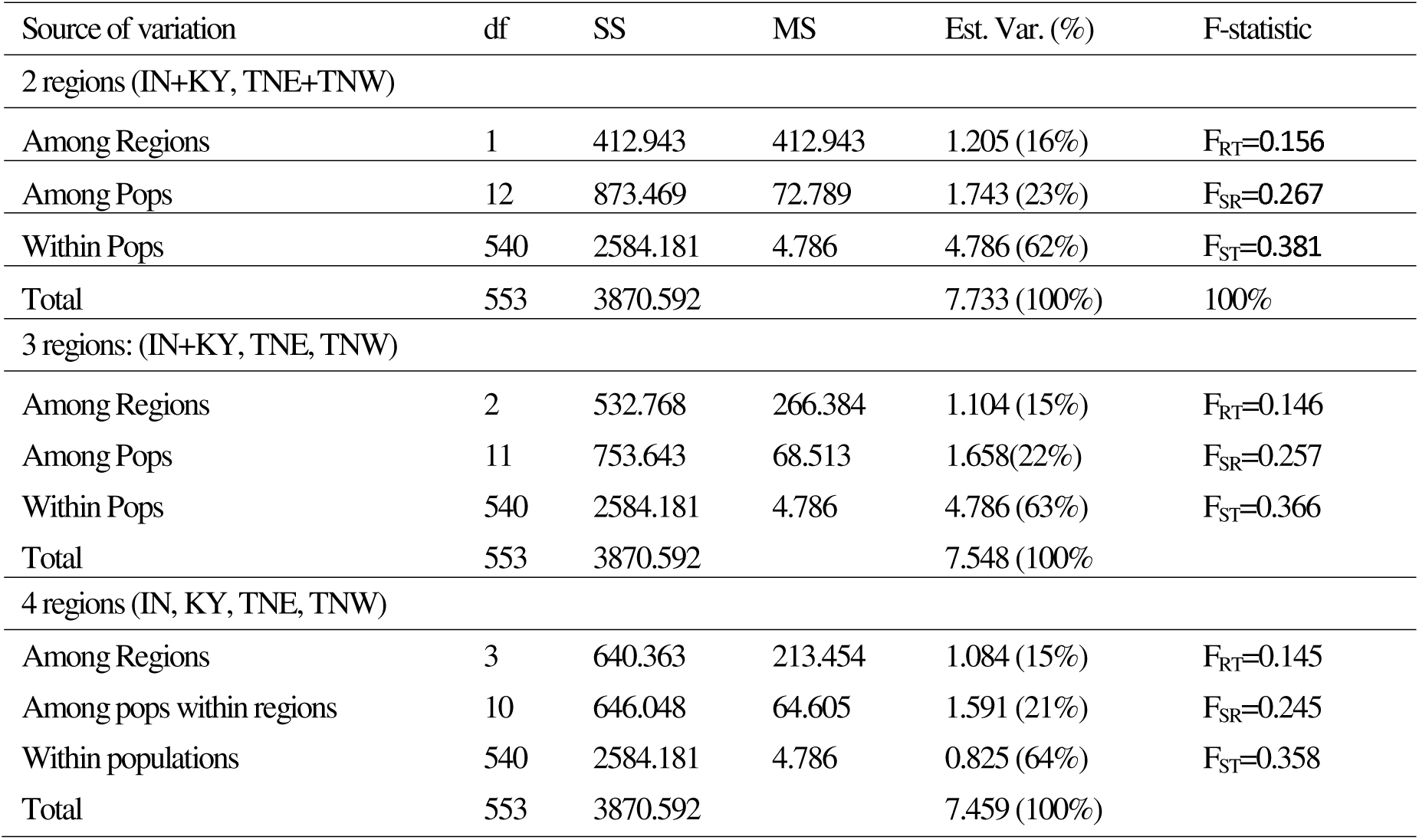
AMOVA summary table with the Indiana (IN), Kentucky (KY), Tennessee east (TNE), and Tennessee west (TNW) populations divided into 1) two groups (IN+KY, TNE+TNW), 2) three groups (IN+KY, TNE, TNW), or 3) the four ecoregions they occupy: Indiana (IN), Kentucky (KY), Tennessee east (TNE), and Tennessee west (TNW),

### Microsatellite markers and genotyping

We developed and employed 20 microsatellite markers to genotype *P. globosa*. We first tested 16 microsatellite markers previously developed for *Physaria filiformis* (C. E. Edwards et al. 2021), of which 10 produced polymorphic PCR bands of the expected size in *P. globosa*. We then developed 10 additional microsatellite markers for *P. globosa* using the protocol described in Swift, Smith, Menges, Bassuner, and Edwards (2016). Briefly, whole-genome libraries were prepared for two *P. globosa* individuals, one from Tennessee and one from Kentucky, using Nextera DNA Sample Prep Kits. Libraries were sequenced on an Illumina MiSeq platform with 2 × 150 bp reads. Sequencing yielded 1,049,048 and 1,163,933 paired-end reads, respectively, which were assembled de novo into 585,371 contigs using the medium sensitivity/fast setting in Geneious v.7.1.9 (Biomatters, Auckland, New Zealand). MSATCOMMANDER (Faircloth 2008) identified 2,054 unique microsatellites from the contig consensus sequences, and PRIMER3 (Rozen & Skaletsky 1999) identified 324 unique primer pairs. An M13 tag (5⍰-CACGACGTTGTAAAACGAC-3⍰) was added to the 5lil end of each forward primer to enable universal dye labeling (Boutin-Ganache, Raposo, Raymond, & Deschepper 2001). Of 48 primer pairs tested, 10 additional polymorphic loci showed reliable amplification in *P. globosa*. These 20 loci (Appendix 1, Table 1) were used to genotype the 406 samples.

PCR amplifications were performed in 10 μL reactions containing 0.5 U GoTaq Flexi DNA polymerase (Promega, Madison, WI, USA), 1× Colorless GoTaq Flexi Buffer (Promega, Madison, WI, USA), 1.5 mM MgCl2, 4.5 pmol each of the reverse primer and one of four fluorescently labeled M13 primers (6-FAM, VIC, NED, or PET; Applied Biosystems, Foster City, CA, USA), 0.18 pmol of the M13-tagged forward primer, and 0.5 mM of each dNTP. Thermal cycling consisted of an initial denaturation at 94 °C for 3 min; 35 cycles of 94 °C for 30 s, 52 °C for 30 s, and 72 °C for 45 s; and a final extension at 72 °C for 20 min. Genotyping was conducted by capillary electrophoresis on an Applied Biosystems 3730xl (Applied Biosystems, Foster City, CA, USA) at the Cornell Biotechnology Resource Center. Raw trace files were scored in GeneMarker v.2.6.2 (SoftGenetics, State College, PA, USA).

### Analysis of population structure

Population genetic structure was characterized by Nei’s standard genetic distance among populations, pairwise F_ST_, and pairwise G_ST_, calculated in Genodive (Meirmans & Van Tienderen 2004). The Nei’s *D* matrix was used to construct a neighbor-joining tree in R using the ape package (Paradis, Claude, & Strimmer 2004), rooted with *P. gracilis* as the outgroup. Genetic structure within *P. globosa* was also examined by PCA in the R package `adegenet` (Jombart 2008) and by Bayesian clustering in STRUCTURE v.2.3.4 (Pritchard, Stephens, & Donnelly 2000). We ran five replicate analyses for each *K* from 1 to 13, using a burn-in of 600,000 generations and a run length of 1,000,000 generations, and evaluated the most likely number of clusters in STRUCTURE HARVESTER (Earl & vonHoldt 2012). The result was visualized on a map using the R package scatterpie (Yu 2025). Hierarchical genetic structure was further quantified by AMOVA in GenAlEx (Peakall & Smouse 2012). We explored three population grouping strategies in AMOVA to determine which best explained patterns of among-population genetic structure, including two groups (IN+KY and TNE and TNW), three groups (IN+KY, TNE, and TNW), and four groups, with each ecoregion in a separate group.

### Analysis of genetic diversity

Samples with >50% missing data were excluded, removing nine *P. globosa* individuals and leaving a final dataset of 397 individuals. The mean number of alleles per population (A), observed and expected heterozygosity (H_O_, H_E_), and the inbreeding coefficient (F_IS_) were calculated by Genodive (Meirmans & Van Tienderen 2004). Private alleles (A_P_) were identified by GenAlEx (Peakall & Smouse 2012), and allelic richness (A_R_) was estimated by HP-Rare (Kalinowski 2005). Null allele frequency (N), genotyping error (b), and the inbreeding coefficient corrected for null alleles (F_B_) were estimated by INEST v.2.2 (Chybicki & Burczyk 2009) under a Bayesian MCMC framework with 5,000,000 cycles, sampling every 250 cycles after a burn-in of 500,000. Recent bottlenecks were assessed by the bottleneck function in INEST v.2.2 based on heterozygosity excess and M-ratio deficiency, each evaluated with a Z-test and a Wilcoxon signed-rank test. Populations were considered to show evidence of a recent bottleneck when at least three of the four tests were significant (*p* < 0.05).

## Results

### Interspecific and intraspecific differentiations

We sampled *P. globosa* from one natural population in the Indiana region (IN), six natural populations in the Kentucky region (KY), five natural populations in western Tennessee (TNW), two natural populations in eastern Tennessee (TNE), and two translocated populations **(Fig. 1A, B**; **Fig. 2).** Broad-scale population structure was consistent across the neighbor-joining tree, STRUCTURE, PCA, and pairwise differentiation analyses. The neighbor-joining tree **(Fig. 1C)**, rooted with *P. gracilis*, placed the three *P. filiformis* populations in a group with *P. gracilis*, while the three *P. ouachitensis* populations formed a sister group to *P. globosa*. Within *P. globosa*, populations were divided into 2 major groups: an IN–KY group and a TNE–TNW group. Pairwise F_ST_ and G_ST_ showed a similar pattern **(Fig. 3C)**, further supporting this major split among natural populations. Consistent with this result, PCA separated IN–KY populations from TNE–TNW populations along PC1, which explained 16.7% of the variance **(Fig. 3B)**.

**Figure 3.**
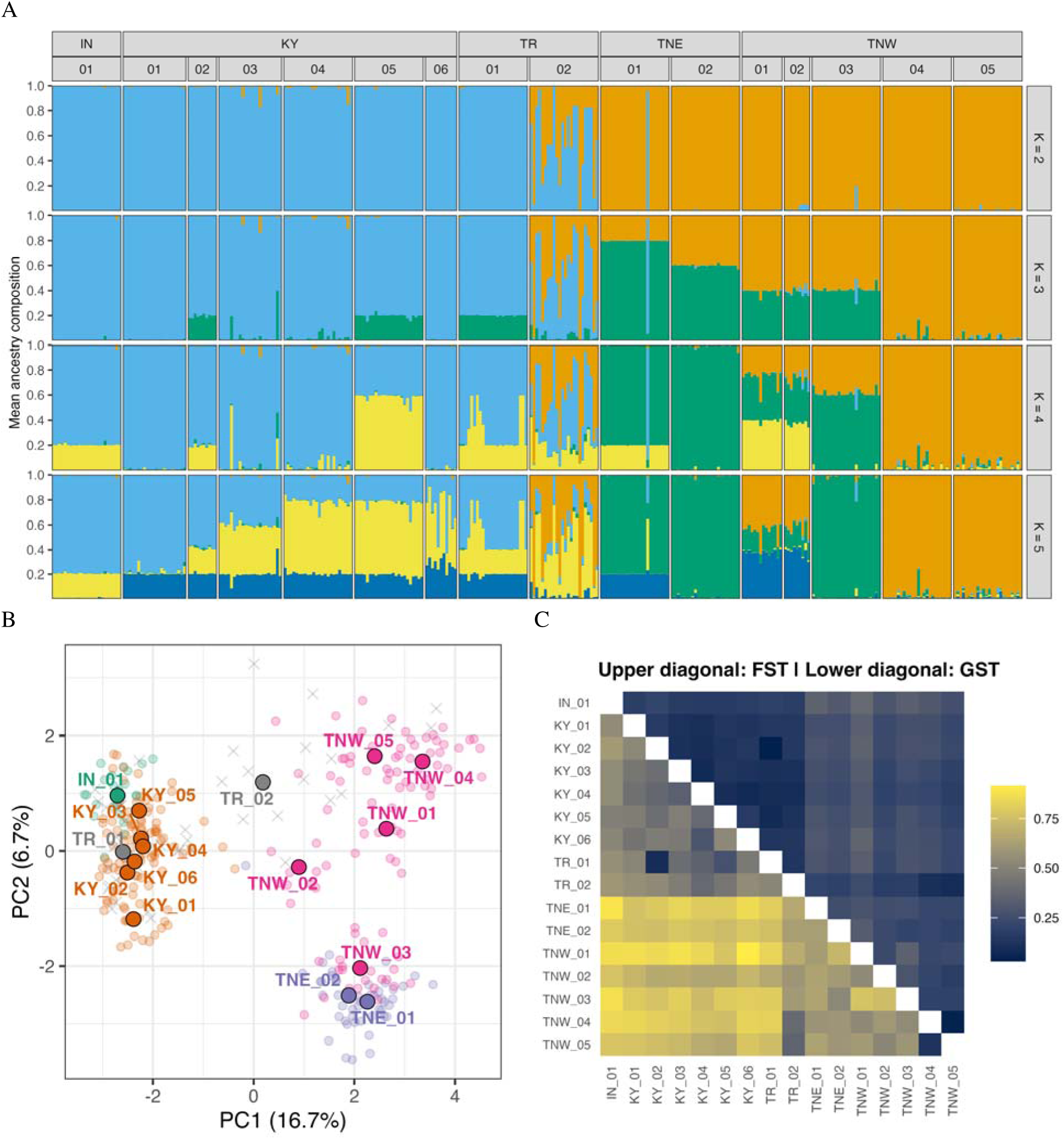
PCA of all natural *P. globosa* populations, with location and species indicated. (A) STURCTURE analysis result (K = 2-5). (B) PCA plot. The big points with black stroke outline are population means. The small points without black stroke outline are individuals from natural populations. The crosses are individuals from *ex situ* populations. Points are colored by regions and population codes are labeled. (C) heatmap of F_ST_ and G_ST_. See **Table 1** for the population codes.

STRUCTURE analyses provided additional resolution of this pattern. Across analyses of *K* = 1–13, the Evanno method identified *K* = 2 as the best-supported number of clusters. At *K* = 2, the IN and KY populations formed a genetic cluster and were separated from the 2^nd^ genetic cluster composed of TNE and TNW populations (**Fig. 3A**). Exploration of higher *K* values revealed finer-scale structure. At *K* = 3, a gradient-like pattern was evident among the TNE and TNW populations, except for the TNW_04 and TNW_05, which remained genetically distinct and largely unadmixed through the highest *K* examined. The Nashville (TN) cosmopolitan area did not block the natural gene flow between the TNE and TNW populations (**Fig. 2**). Instead, the genetic distinctiveness of the TNW_04 and TNW_05 populations could be a result of elevated inbreeding (**Fig. 4D**). At *K* = 5, the IN population formed its own genetic cluster, although it remained more similar to the KY than to the TNE and TNW populations, and a gradient-like pattern was also visible among the KY populations (**Fig. 3A**). PCA supported these finer-scale results: PC2, which explained 6.7% of the variance, further separated TNE and TNW populations, except for TNW_03, which clustered with the TNE populations (**Fig. 3B**).

**Figure 4.**
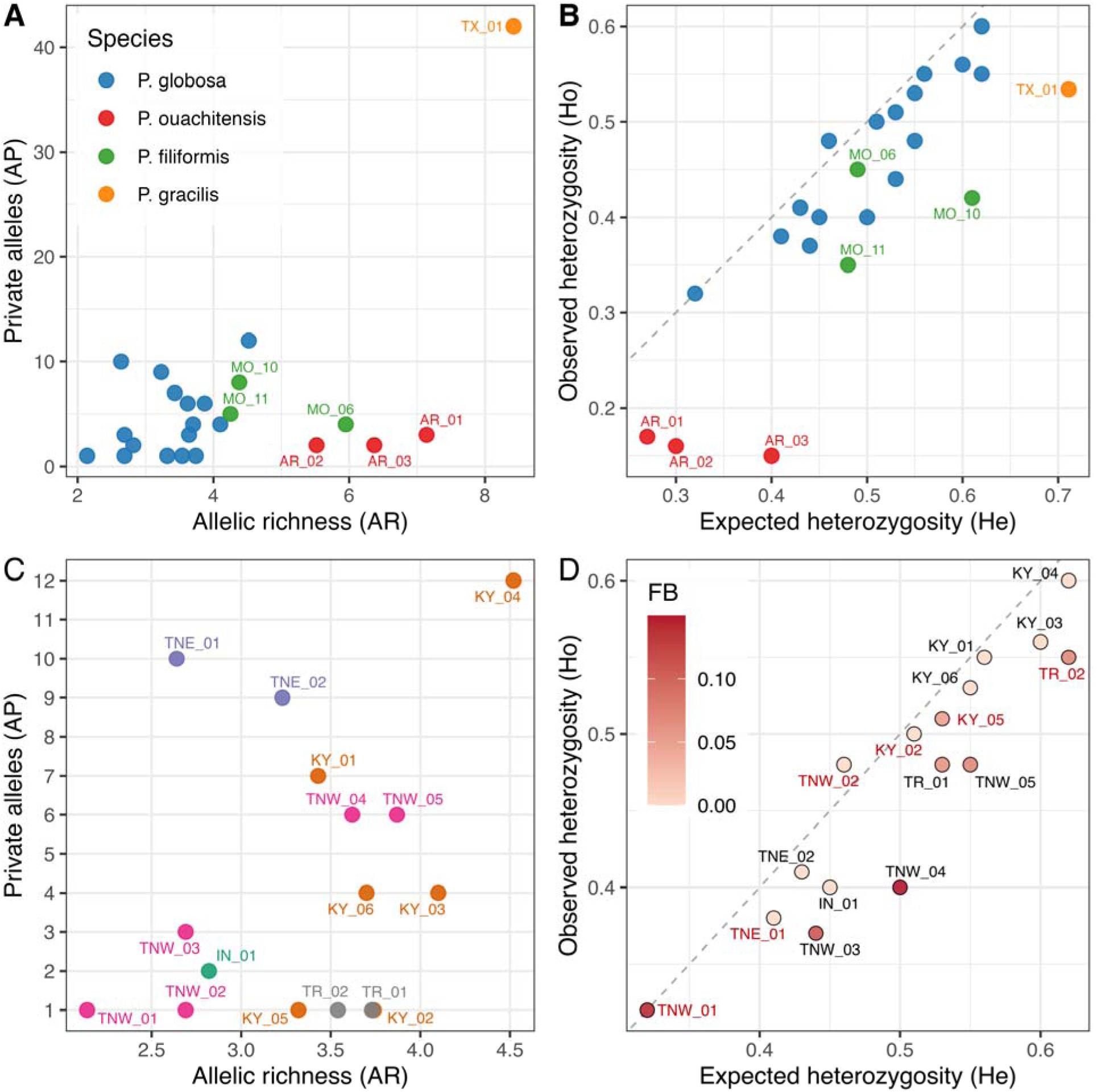
Genetic diversity analyses plots. (**A-B**), *Physaria globosa* and outgroup species. (**C-D**), *Physaria globosa* with population labels. F_B_., inbreeding coefficient after correction for null alleles. The six populations with excessive heterozygosity are labeled in red in (**D**). See **Table 1** for the population codes.

The translocated populations also showed genetic patterns consistent with their documented origins. The TR_01 clustered with the KY populations in both STRUCTURE and PCA, consistent with its origin from the KY_02 and KY_05 populations (**Fig. 3A, B**). At K = 4-5, four individuals (16.7%) of the TR_01 population showed potential admixture between the two source populations, 16 individuals (66.7%) grouped with KY_02 and four individuals (16.7%) grouped with KY_05 (**Fig. 3A**). The KY_02 population had more genetic contributions to the TR_01 population than the KY_05 population (**Fig. 3C**). The TR_02 population was intermediate between the IN-KY and TNE-TNW populations in PCA and showed admixed ancestry in STRUCTURE (**Fig. 3A, B**). At *K* = 5, STRUCTURE indicated that the TR_02 population included individuals with varying degrees of admixture, including putative F_1_ hybrids and backcrosses (**Fig. 3A**). In addition, G_ST_ indicated that the TR_02 population received stronger genetic contributions from the TNW_04 and TNW_05 populations than from other Tennessee populations (**Fig. 3C**).

Among natural populations, we detected significant isolation by geographic distance (IBD) (Mantel’s *r* = 0.505, *p* = 0.001). AMOVA further showed that genetic variation was partitioned across multiple hierarchical levels, with the partitioning of genetic variation similar regardless of whether populations were grouped into two, three, or four groups (**Table 2**). When populations were divided into two groups, 16% of the variation occurred between regions (IN+KY, TNE, and TNW), 23% among populations within regions, and 62% within populations. When populations were divided into three groups, 15% of the variation occurred between regions (IN+KY, TNE+TNW), 22% among populations within regions, and 63 % within populations.When populations were grouped into four groups, 15% of the variation occurred among regions (IN, KY, TNE, and TNW), 21% among populations within regions, and 64% within populations. The fact that two groups explained the greatest among-population variation is in agreement with the results of analyses of PCA and STRUCTURE.

### Genetic diversity, inbreeding, and bottleneck

Populations of *P. globosa* varied in their genetic diversity (**Table 3**). The average number of alleles per locus (A) in *P. globosa* ranged from 2.278 – 6.5, the number of private alleles per population (A_P_) ranged from 1-12, and the allelic richness (A_R_) ranged from 2.14 – 4.52 (**Table 3**). *P. globosa* had slightly lower A_R_ than its rare sister species, *P. filiformis and P. ouachitensis*, and much lower A_R_ than the widespread species, *P. gracilis*. The *P. gracilis* population had an extraordinarily high number of private alleles (A_P_) because we only sampled from one population. Within *P. globosa*, both private alleles per population (A_P_) and the allele richness (A_R_) were lowest in the population TNW_01 from western Tennessee (TNW), and greatest in population KY_04 from Kentucky (KY) (Fig. 4). The translocated (TR) populations had intermediate A_R_ but low A_P_, which is consistent with genetic mixing of two or more populations.

**Table 3.** Genetic diversity metrics for *Physaria globosa* and related species.

| Population | Region | n | A | A<sub>R</sub> | A<sub>P</sub> | H<sub>O</sub> | H<sub>E</sub> | F<sub>IS</sub> | % null | F<sup>B</sup> |
| --- | --- | --- | --- | --- | --- | --- | --- | --- | --- | --- |
| *P. globosa* |  |  |  |  |  |  |  |  |  |  |
| TR_01 | KY | 24 | 4.72 | 3.73 | 1 | 0.48 | 0.53 | 0.08 | 0.46 | 0.06 |
| TR_02 | KY | 24 | 4.39 | 3.54 | 1 | 0.55 | 0.62 | 0.09 | 0.18 | 0.07 |
| IN_01 | IN | 24 | 3.28 | 2.82 | 2 | 0.4 | 0.45 | 0.08 | 0.16 | 0 |
| KY_01 | KY | 22 | 4.11 | 3.43 | 7 | 0.55 | 0.56 | 0 | 0.22 | 0 |
| KY_02 | KY | 10 | 4 | 3.74 | 1 | 0.5 | 0.51 | 0.05 | 0.18 | 0 |
| KY_03 | KY | 20 | 5.33 | 4.1 | 4 | 0.56 | 0.6 | 0.07 | 0.3 | 0 |
| KY_04 | KY | 24 | 6.5 | 4.52 | 12 | 0.6 | 0.62 | 0.03 | 0.26 | 0 |
| KY_05 | KY | 24 | 3.94 | 3.32 | 1 | 0.51 | 0.53 | 0.03 | 0.18 | 0.05 |
| KY_06 | KY | 11 | 4.06 | 3.7 | 4 | 0.53 | 0.55 | 0.06 | 0.28 | 0 |
| TNE_01 | TNE | 23 | 3.22 | 2.64 | 10 | 0.38 | 0.41 | 0.11 | 0.15 | 0 |
| TNE_02 | TNE | 24 | 4.17 | 3.23 | 9 | 0.41 | 0.43 | 0.02 | 0.33 | 0 |
| TNW_01 | TNW | 14 | 2.28 | 2.14 | 1 | 0.32 | 0.32 | 0.03 | 0.08 | 0.14 |
| TNW_02 | TNW | 9 | 2.78 | 2.69 | 1 | 0.48 | 0.46 | -0.04 | 0.17 | 0 |
| TNW_03 | TNW | 24 | 3.28 | 2.69 | 3 | 0.37 | 0.44 | 0.23 | 0.47 | 0.11 |
| TNW_04 | TNW | 24 | 4.72 | 3.62 | 6 | 0.4 | 0.5 | 0.16 | 0.33 | 0.15 |
| TNW_05 | TNW | 24 | 5.44 | 3.87 | 6 | 0.48 | 0.55 | 0.11 | 0.36 | 0.07 |
| Average across *P. globosa* |  |  | 4.2 | 3.38 | 4.24 | 0.47 | 0.51 | 0.08 | 0.26 | 0.04 |
| *P. ouachitensis* |  |  |  |  |  |  |  |  |  |  |
| AR_01 | AR | 5 | 1.79 | 7.14 | 3 | 0.17 | 0.27 | 0.37 | 0.45 | 0 |
| AR_02 | AR | 9 | 2.4 | 5.52 | 2 | 0.16 | 0.3 | 0.46 | 0.53 | 0.38 |
| AR_03 | AR | 4 | 1.85 | 6.37 | 2 | 0.15 | 0.4 | 0.61 | 0.57 | 0 |
| Average across *P. ouachitensis* |  |  | 2.01 | 6.34 | 2.33 | 0.16 | 0.32 | 0.48 | 0.52 | 0.13 |
| *P. filiformis* |  |  |  |  |  |  |  |  |  |  |
| MO_06 | MO | 4 | 2.53 | 5.95 | 4 | 0.45 | 0.49 | 0.08 | 0.24 | 0 |
| MO_10 | MO | 6 | 3.18 | 4.38 | 8 | 0.42 | 0.61 | 0.32 | 0.54 | 0 |
| MO_11 | MO | 7 | 2.81 | 4.25 | 5 | 0.35 | 0.48 | 0.29 | 0.42 | 0 |
| Average across *P. filiformis* |  |  | 2.84 | 4.86 | 5.67 | 0.4 | 0.53 | 0.23 | 0.4 | 0 |
| *P. gracilis* |  |  |  |  |  |  |  |  |  |  |
| TX_01 | TX | 11 | 5 | 8.42 | 42 | 0.534 | 0.711 | 0.249 | 0.77 | 0 |

Column definitions: n=number of individuals; A=average alleles per locus; A<sub>R</sub>=allelic richness; A<sub>P</sub>=private alleles per population; H<sub>O</sub>=observed heterozygosity; H<sub>E</sub>=expected heterozygosity; F=fixation index; % null=percentage of null alleles; F<sub>B</sub>=fixation index adjusted for null alleles.

In *Physaria globosa,* observed heterozygosity (H_O_) ranged from 0.323 – 0.596, and expected heterozygosity (H_E_) ranged from 0.323 – 0.620, which is similar to *P. filiformis* and *P. gracilis*, but greater than *P. ouachitensis.* Within *P. globosa*, most populations from Kentucky (KY) exhibited greater than average H_O_ and H_E_, and populations from both eastern Tennessee (TNE) and western Tennessee (TNW) exhibited lower than average H_O_ and H_E_ (**Table 3**). Generally, H_E_ was slightly greater than H_O_ in most populations, such that the average inbreeding coefficient (F) was slightly positive (overall mean = 0.08, range -0.036 to 0.227) (**Table 3**). However, these positive inbreeding coefficients are likely due in part to null alleles rather than inbreeding, as the INEST analysis revealed that the percentage of null alleles (% null) ranged from 0.0771 to 0.4694 (overall mean = 0.2648) (Table 3). After correcting for null alleles, inbreeding coefficients (F_B_) were generally very close to zero (range 0 – 0.1481; overall mean = 0.04; Table 3), indicating that most populations have not experienced inbreeding. However, three populations (TNW_04, TNW_03, and TNW_01), all from the TNW region, exhibited inbreeding coefficients greater than >0.10 even when corrected for null alleles, suggesting that they may have experienced some inbreeding.

We used HP-rare to test for genetic bottlenecks and considered populations to show significant evidence for a genetic bottleneck when three out of the four tests were statistically significant. This revealed that the following six populations, representing three geographic regions, had undergone recent bottlenecks: populations KY_02, KY_05, TR_02 from the KY region; population TNE_01 from the TNE region; and populations TNW_01 and TNW_02 from TNW region (**Fig. 4**). Generally, populations with the lowest A_R_ also exhibited significant evidence of either genetic bottlenecks (TNE_01, TNW_01, TNW_02), inbreeding (TNW_01 and TNW_03), or both (TNW_01).

## Discussion

Our results indicate that conservation planning for *Physaria globosa* must balance two genetic realities: the species retains strong regional genetic structure that warrants conservation, but genetic vulnerability is unevenly distributed and may require population-specific interventions. *Physaria globosa* is neither genetically homogeneous across its range nor uniformly genetically depauperate across populations. Instead, the species exhibits strong regional differentiation and substantial variation in within-population diversity, with some populations displaying extremely low levels of genetic diversity and recent bottlenecks. The translocated populations exhibit clear genetic mixing, indicating that mixing material from different source populations can successfully produce viable offspring and increase genetic variation. However, the fitness consequences of genetic mixing remain unknown for this species and should be carefully evaluated. Taken together, these findings suggest that a multi-pronged conservation strategy involving protection of natural regional genetic structure, *ex situ* conservation of remaining genetic variation, and mixed-source translocations may help maintain genetic variation in *P. globosa*.

### Genetic diversity and structure in *P. globosa*

*Physaria globosa* exhibits strong genetic structure across its disjunct geographic range, a pattern similar to that reported for the closely related species *Physaria filiformis* (C. E. Edwards et al. 2021). Neighbor-joining analysis, PCA, STRUCTURE, and pairwise genetic differentiation consistently support a major split between the IN–KY and TNE–TNW groups, with additional substructure within each group. This pronounced spatial structure is likely reinforced by discrete habitats and limited dispersal. Seed movement in *P. globosa* appears to be largely gravity-mediated, and pollen dispersal may also be spatially restricted by localized pollinator activity involving bees, flies, and other invertebrates (Waters, Hurt, & Krosnick 2025). In addition, facultative self-compatibility in *P. globosa* may promote elevated inbreeding within populations (Powell 2023), further contributing to localized genetic structure. Significant isolation-by-distance and AMOVA results are consistent with this interpretation, indicating that genetic differentiation is structured both among regions and among populations within regions.

However, genetic structure was not fully explained by geographic distance alone. Two Tennessee populations (TNE_02 and TNW_03) were genetically similar despite occurring on opposite sides of the greater Nashville metropolitan area (Fig. 2). Given that these populations both occur along the Cumberland River, their similarity may reflect rare dispersal events via hydrological processes, flooding, animal vectors, human activity, or transportation corridors. For example, historical connectivity via the Tennessee Central Railway may have facilitated low levels of gene flow or movement of propagules (Hicks 2023). Thus, even though *P. globosa* populations are strongly genetically structured overall, occasional dispersal or historical connectivity may contribute to genetic similarity among geographically distant populations.

Genetic diversity in *P. globosa* was unevenly distributed across its range. Allelic richness (A_R_) was generally lower in *P. globosa* than in its congeners, but diversity also varied substantially among populations. Generally, diversity was greater in the northern part of the species’ range in IN and KY, whereas genetic diversity was lower in the TN populations. For example, KY_04 exhibited the greatest allelic richness and number of private alleles, whereas TNW_01 had the lowest. Bottleneck and inbreeding analyses support this pattern, indicating that genetic risk is localized in the southern portion of the species’ range rather than pervasive across the species. Spatial variation in genetic diversity may, in part, reflect differences in disturbance and management histories. In the southern portion of the range, construction of locks and dams along the Cumberland River (Olson 2021), historical railroad development (Hicks 2023), roadside herbicide use, recreation, and development (Long et al. 2021) may have reduced population sizes and altered connectivity in TN. Invasive *Lonicera maackii* may further contribute to local genetic erosion by reducing light availability and decreasing native plant growth (McNeish & McEwan 2016), a threat particularly relevant for a species potentially dependent on high-light conditions maintained by periodic disturbance (Krosnick et al. 2022; Long et al. 2021). Conversely, KY_04, which has been regularly managed with *L. maackii* removal since 2018, exhibited among the highest genetic diversity of all sampled populations. Although our data cannot directly test whether these management practices promoted high diversity, this pattern is consistent with the hypothesis that active habitat management may promote recruitment and help maintain genetic variation, possibly through contributions from a persistent soil seed bank.

### Genetic variation in mixed-source translocations

Genetic mixing of multiple source populations can be used in *ex situ* and translocated populations to reduce inbreeding and increase gene flow and genetic diversity, but such intervention should be guided by regional genetic structure, source-population history, local adaptation, and post-translocation monitoring rather than treated as universally beneficial (Van Rossum et al. 2025; Van Rossum & Le Pajolec 2021). In *Physaria globosa*, this concern is particularly relevant because genetic mixing and translocation efforts have been conducted for decades without prior genetic assessment or long-term genetic monitoring. This study documented the genetic composition of two genetically mixed populations whose genetic sources originated across different geographic scales. One genetically mixed population (TR_01) was sourced from two populations within the same geographic region, where gene flow between the source populations could occur naturally. In contrast, the other genetically mixed population (TR_02) was sourced from two geographically distinct regions, between which natural gene flow rarely occurs.

Translocations that promote genetic mixing and increase genetic diversity may be less controversial when they restore or augment gene flow among populations that are naturally or historically connected, because such interventions are less likely to disrupt historical population structure or local adaptation, if present. The TR_01 population was initially established using seeds from the KY_02 population and was subsequently augmented with seeds from the KY_05 population. Based on our population structure analysis, natural gene flow has occurred between these two source populations. There is no sign of population bottleneck in the TR_01 population, suggesting that the founder population of 100-200 seeds from 30-40 maternal plants was sufficient to maintain a level of genetic diversity comparable to that observed in natural populations. When we sampled the TR_01 population in 2021, it had been 7 years since the introduction of the KY_02 population and 6 years since the introduction of the KY_05 population. However, we only detected that 16.7% of the population was admixed. The low proportion of admixture could be attributable in part to the facultative self-compatibility exhibited by the species. However, it is likely mostly attributable to the mostly biennial life history of the species. Given that more than 95% of *P. globosa* individuals exhibited a biennial life history, flowering in their 2^nd^ year in a greenhouse experiment (unpublished data), individuals sourced from KY_02 and KY_05 and introduced in different years may have had limited overlap in flowering phenology, thereby reducing opportunities for genetic mixing. These findings suggest that flowering phenology may need to be taken into consideration for managing reintroductions of biennial species. Specifically, if mixing of different source populations is desired, then they may need to be introduced in the same year, or new seed could be introduced in the same year that existing plants are flowering.

In contrast, although translocations that mix naturally isolated populations can restore genetic diversity and alleviate inbreeding depression, they may also alter natural population structure, demographic history, expression of genetic load, or local adaptation, if present (Bell et al. 2019; Frankham 2015; Pérez-Pereira, Caballero, & García-Dorado 2022; Sexton, Strauss, & Rice 2011). Such interventions should therefore be considered only after careful evaluation and when the target species faces a high risk of extinction without intervention. Our genetic analysis indicated that the TR_02 population is a mixture of genotypes from the IN–KY and TNE–TNW regions, which are geographically distinct and experience little natural gene flow. Although this result demonstrates that individuals from these geographically distinct regions can produce viable offspring, for the reasons discussed above, *we do not recommend mixing source populations from different regions without first evaluating offspring performance and genetic compatibility in a controlled setting*. Moreover, this translocated population also showed a significant recent bottleneck with a deficit of rare alleles at segregation sites, which implies this population was founded with few individuals instead of through well-planned genetic management. Importantly, although some *P. globosa* populations have experienced demographic bottlenecks and reduced genetic diversity, substantial genetic diversity is still maintained within geographic regions, and there are no clear signs that the species faces imminent extinction.

### Conservation recommendations

Our genetic findings support a risk-benefit framework for conservation management for *P. globosa*. Strong regional structure (IN-KY vs. TNE-TNW) argues against treating all populations as interchangeable seed sources, as such practices could erode historic genetic differentiation. Conversely, reduced diversity, bottleneck signals, localized inbreeding, and ongoing habitat threats suggest that current management practices alone may be insufficient in some cases. The most defensible interpretation is therefore not that management must choose between conserving population structure or restoring gene flow, but that both goals must be balanced, such that a combination of in situ and ex situ conservation may be optimal.

*In situ* conservation should remain the primary focus because it maintains the geographic distribution of genetic diversity and supports representation and redundancy across the range. Efforts should prioritize the protection and management of multiple populations within each of the major geographic regions (IN, KY, TNE, TNW). At present, Indiana (IN) may be the only region where this goal is largely met, because its single population is both protected and actively managed. In eastern Tennessee (TNE), several populations along the Cumberland River are protected, and populations associated with cliff-faces and ledges may be less vulnerable to invasion by *L. maackii*; however, the genetic status of many of the sites in this region remains uncertain due to limited access during this study. Many populations in Kentucky (KY) and western Tennessee (TNW) occur on private land and are not currently being actively managed, making them important targets for future conservation. Populations showing low allelic richness, bottleneck signatures, elevated inbreeding, or severe habitat degradation should therefore be prioritized for closer demographic and genetic monitoring, habitat management, and, where appropriate, demographic or genetic augmentation. However, these efforts may be logistically challenging because *P. globosa* often occurs on cliff tops, ledges, and steep slopes that are difficult and potentially dangerous to access, making them important targets for *ex situ* conservation.

Where in situ conservation is difficult, e*x situ* conservation and translocations may be complementary approaches to conserve *P. globosa*. Whenever possible, we recommend that these efforts be guided by genetic data to ensure that only populations that have experienced recent gene flow are mixed and that these efforts are coupled with continued monitoring and evaluation to assess their success. One example of where this could be useful is in the population TNW_01, which showed the lowest allelic richness and heterozygosity among all studied populations. If future studies show that TNW_01 has experienced rapid population decline and strong inbreeding depression, we would suggest conducting genetic rescue by translocating individuals from TNW_04 or TNW_05, which are closely related but have higher genetic diversity, into TNW_01.

Finally, this study highlights the limitation of inference based solely on neutral microsatellite markers. Although these markers effectively characterize genetic structure, relative population genetic diversity, and admixture, they do not provide insight into adaptive variation among regional groups or the fitness consequences of genetic mixing for translocation. This uncertainty is particularly relevant for a species in which conservation decisions may involve translocating genotypes among historically differentiated populations. Future work integrating demographic data, common-garden or fitness experiments, and genome-scale markers will be necessary to determine whether management interventions and multi-source conservation translocations enhance long-term persistence and evolutionary potential in *P. globosa*.

## Acknowledgements

We thank Geoff Call, Caitlin Elam, Noah Dell, Julian Campbell, Shawn Krosnick, Scott Namestnik, and John Malone for assistance with funding, collection permits, and sampling.

## Statements & Declarations

### Author Contributions

W.-H. K. analyzed the data, visualized the results, and wrote the manuscript. C. L. analyzed the data and wrote the manuscript. B. B. conducted molecular experiments. A. G. L. conducted fieldwork, collected samples, and revised the manuscript. T. R. L. conducted fieldwork, collected samples, and wrote the manuscript. M. A. A. conceived the project, conducted fieldwork, collected samples, and revised the manuscript. C. E. E. conceived of the project, obtained funding, q ap0-1q2conducted fieldwork, collected samples, analyzed the data, and wrote the manuscript. All authors read and approved the final manuscript.

## Data availability

The datasets generated and analyzed during the current study are available in the Physaria_SSR repository, https://github.com/kuowenhsi/Physaria_SSR.

